# Familial migration of the Neolithic contrasts massive male migration during Bronze Age in Europe inferred from ancient X chromosomes

**DOI:** 10.1101/078360

**Authors:** Amy Goldberg, Torsten Günther, Noah A. Rosenberg, Mattias Jakobsson

**Affiliations:** Department of Biology, Stanford University, 94305 Stanford, California; Department of Organismal Biology, Uppsala University, 75236 Uppsala, Sweden; Science for Life laboratory, Uppsala University, 75123 Uppsala, Sweden

## Abstract

Dramatic events in human prehistory, such as the spread of agriculture to Europe from Anatolia and the Late Neolithic/Bronze Age (LNBA) migration from the Pontic-Caspian steppe, can be investigated using patterns of genetic variation among the people that lived in those times. In particular, studies of differing female and male demographic histories on the basis of ancient genomes can provide information about complexities of social structures and cultural interactions in prehistoric populations. We use a mechanistic admixture model to compare the sex-specifically-inherited X chromosome to the autosomes in 20 early Neolithic and 16 LNBA human remains. Contrary to previous hypotheses suggested by the patrilocality of many agricultural populations, we find no evidence of sex-biased admixture during the migration that spread farming across Europe during the early Neolithic. For later migrations from the Pontic steppe during the LNBA, however, we estimate a dramatic male bias, with ~5-14 migrating males for every migrating female. We find evidence of ongoing, primarily male, migration from the steppe to central Europe over a period of multiple generations, with a level of sex bias that excludes a pulse migration during a single generation. The contrasting patterns of sex-specific migration during these two migrations suggest a view of differing cultural histories in which the Neolithic transition was driven by mass migration of both males and females in roughly equal numbers, perhaps whole families, whereas the later Bronze Age migration and cultural shift were instead driven by male migration, potentially connected to new technology and conquest.

---

Genetic data suggest that modern European ancestry represents a mosaic of ancestral contributions from multiple waves of prehistoric migration events. Recent studies of genomic variation in prehistoric human remains have demonstrated that two mass migration events are particularly important to understanding European prehistory: the Neolithic spread of agriculture from Anatolia starting ~9,000 years ago, and migration from the Pontic-Caspian Steppe ~5,000 years ago (1–6). These migrations are coincident with large social, cultural, and linguistic changes, and each has been inferred to replace more than half the gene pool of resident Central Europeans during those times.

During such events, males and females often experience different demographic histories owing to cultural factors such as norms regarding inheritance and the residence locations of families in relation to parental residence, social hierarchy, sex-biased admixture, and inbreeding avoidance (7–11). Empirical evidence suggests that sex-specific differences in migration and admixture have shaped patterns of human genomic variation worldwide, with notable examples occurring in Africa, Austronesia, Central Asia, and the Americas (12–15). These sex-specific behaviors leave signatures in the patterns of variation in genetic material that is differentially inherited between males and females in a population. Therefore, contrasting patterns of genetic variation for differentially inherited genetic material can be informative about past sociocultural and demographic events (7–11,16).

Analyses of the maternally inherited mitochondrial DNA (mtDNA) and the paternally inherited Y chromosome have lent differential support for the hypothesis that the Neolithic spread of agriculture from Anatolia occurred through a large population migration, rather than a spread of technology (17–21). In general, studies of Y-chromosomal data more than mtDNA have supported Anatolian migration, which has been interpreted as evidence for male-biased migration of the population that introduced farming. This hypothesis of male-biased migration of farming populations is consistent with ethnographic studies showing a higher frequency of patrilocality in farming than hunter-gatherer populations, as an inheritance model through the paternal lineage would favor the persistence of farming-associated Y chromosomes with more flexibility for the source population of female mates. Isotopic studies from Neolithic European archeological sites suggest more female than male migration on a local scale, supporting the shift to patrilocality in the region (9,22).

Based on archeological and modern-day genetic data, the later migration from the Pontic-Caspian Steppe has also been hypothesized to be male-biased (23–25). Populations in the region, such as the Yamnaya or Pit Grave culture, are thought to have strong male-biased hierarchy, as inferred by overrepresentation of male burials, male deities, and kinship terms (26). The region is a putative origin for the domesticated horse in Europe, and the culture is known for its use of horse-driven chariots, a potential male-biased mechanism of dispersal into central Europe (26).

Combining recent analytical advances in the understanding of admixture on the autosomes and the sex-specifically inherited X chromosome with technological advances that have generated genome-wide data from many ancient samples now makes it possible to consider the contrasting male and female genetic histories of prehistoric Europe. We test the hypotheses that migrations from Anatolia during the Neolithic Transition and from the Pontic Steppe during the late Neolithic/Bronze Age period were male-biased.

Figure 1 provides a schematic of the population admixture events that have previously been inferred (1–6). Previous studies have inferred the relationship between populations, but did not consider a population history model. We compare genetic differentiation of the autosomes and the X chromosome between the migrating and admixed populations for each migration event: Anatolian farmers (AF) to early Neolithic Central Europeans (CE) and Pontic Steppe pastoralists (SP) to late Neolithic and Bronze Age Central Europeans (BA). We compute the statistic *Q* (27,28), which is an estimator of the ratio of effective population size of the X chromosome to that of the autosomes based on *F*_*ST*_ (Materials and Methods). Under a demographic model with equal male and female effective sizes, *Q* is expected to be ¾, as there are three X chromosome for every four autosomes in the population. Deviations from ¾ would therefore show sex-biased effective population sizes, which indicate different population histories for males and females. Comparing AF and CE populations for the Neolithic transition, X and autosomal differentiation is similar to that expected for a non sex-biased process (Table 1). In contrast, there is high relative differentiation on the X chromosome between SP and BA populations (*Q* = 0.237, Table 1), indicating strong male bias during the Pontic steppe migration.

**Fig. 1:**
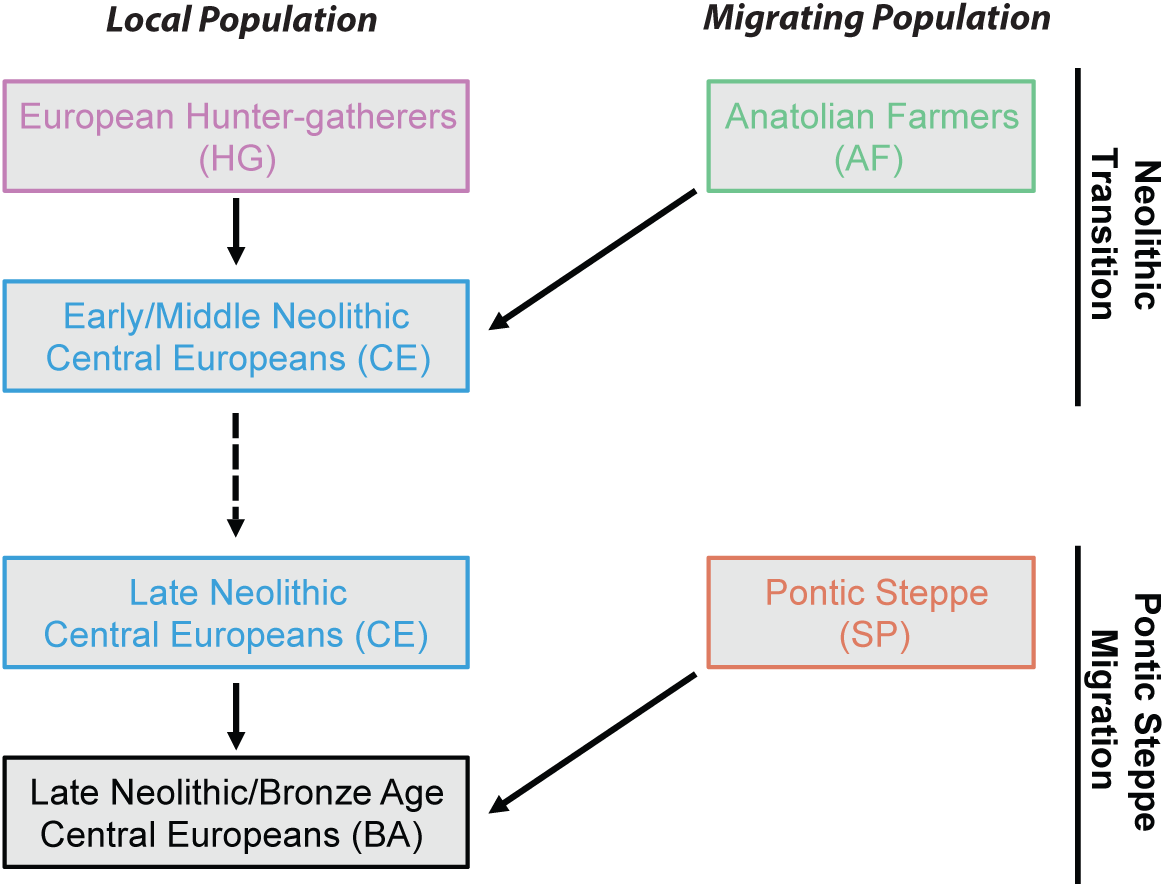
Schematic of the admixture history of Central European farmers during the Neolithic and Bronze Age. First, a migration from Anatolia occurred during the Neolithic transition, and second, a late Neolithic/Bronze Age migration occurred from the Pontic-Caspian Steppe to central Europe. In both cases, the migrating population mixed with the contemporaneous local population upon entering central Europe.

**Table 1:** Comparisons of *F*_*ST*_ on the X chromosome and autosomes. The quantity *Q* compares genetic differentiation, calculated from *F*_*ST*_, on the X chromosome and autosomes. For an ideal population with no changes in effective size, *Q* is expected to be ¾ (Materials and Methods). Notably, *Q* is close to ¾ for the CE-AF comparisons, but is considerably lower for the BA-SP comparisons.

| Populations | $F_{ST}^A$ | $F_{ST}^X$ | Q |
| --- | --- | --- | --- |
| CE-AF | 0.00352 | 0.00503 | 0.700 |
| CE-HG | 0.0526 | 0.0574 | 0.911 |
| BA-SP | 0.00777 | 0.0319 | 0.237 |
| BA-AF | 0.0250 | 0.0302 | 0.826 |
| BA-HG | 0.0307 | 0.0351 | 0.872 |

In order to infer sex-specific admixture rates and compare potential migration models, we estimated ancestry proportions on the X chromosome and autosomes separately, with a model-based clustering algorithm (29), using the ancient genomes as proxies for the ancient source groups in our population model and employing supervised clustering (Materials & Methods, Fig. 2A, Tables S1-3). For an admixture process with equally many males and females contributing, the ratio of mean X-chromosomal admixture to mean autosomal admixture is expected to be 1. An admixture process with more contributing males leads to a reduction of the migrating population’s ancestry on the X chromosome compared to the autosomes.

**Fig. 2:**
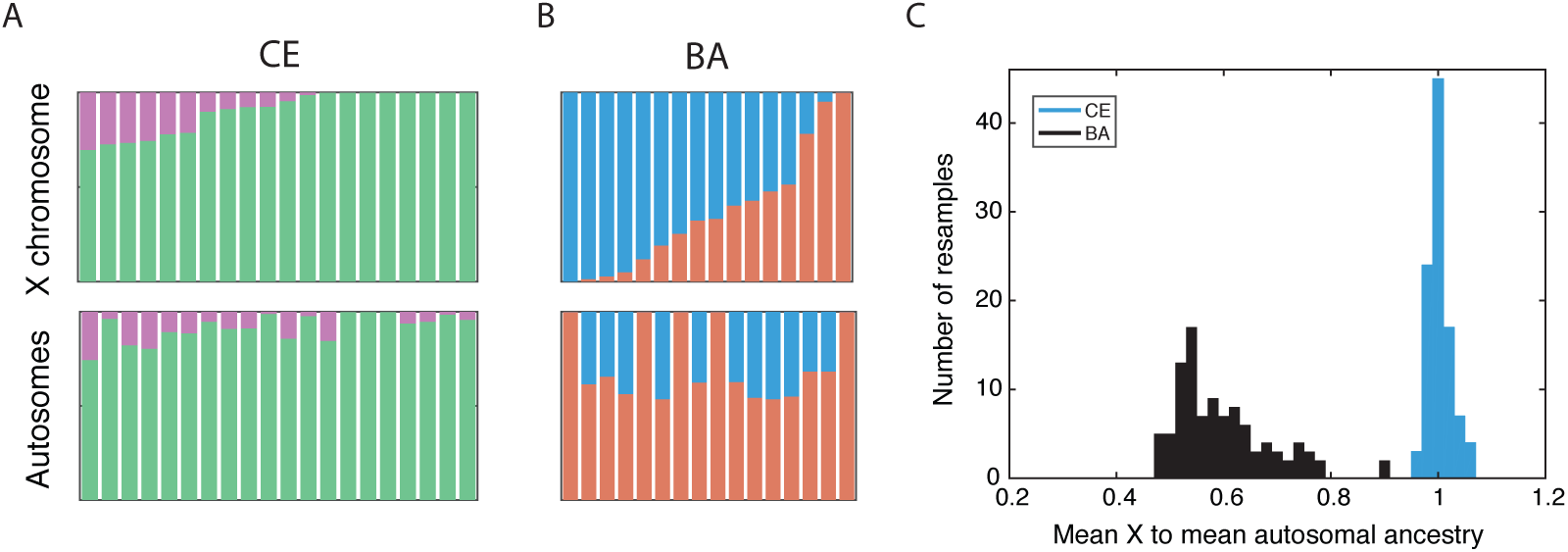
Comparisons of estimated X and autosomal ancestry on the basis of model-based supervised clustering. (A) Early/middle Neolithic Europeans (CE). (B) Late Neolithic/Bronze Age Europeans (BA). Individuals are ordered by X-chromosomal ancestry, with corresponding autosomal ancestry for the same individual shown below. Clustering results by individual are presented in Table S1. (C) Histograms of the ratio of the mean across individuals of X-chromosomal ancestry to the mean across individuals of autosomal ancestry for 100 autosomal resampled estimates using random draws of SNPs equal to the number of X-chromosomal SNPs for the corresponding population (Materials and Methods). Colors for all panels correspond to ancestry groups given in Figure 1

For the Neolithic transition, we estimated the ratio of mean (across individuals) AF ancestry on the X chromosome to the mean on the autosomes as 0.903/0.913 = 0.989, and the corresponding ratio for HG ancestry is 0.097/0.087 = 1.115. Comparing the mean X-chromosomal AF ancestry to the mean autosomal AF ancestry in each of the 100 estimates from resampled autosomal SNPs (Materials and Methods), the median ratio of X to autosomal AF ancestries is 1.00 (Fig. 2B). The mean X-chromosomal admixture ± one standard error estimated by bootstrapping the admixture estimates in 100 resamples of blocks of SNPs largely overlaps with the distribution of mean autosomal ancestry in the population over the 100 estimates (Fig. S1). The distributions of X and autosomal ancestry within the sampled population are not significantly different (*p* = 0.493, Wilcoxon signed-rank test, Fig. S2).

We additionally considered the fraction of individuals in the admixed population with higher X-chromosomal than autosomal ancestry. This measure is indicative of sex bias, with less emphasis on the exact value of the ancestry proportions. Excluding three individuals with 100% ancestry estimated to be from Anatolian-related populations on both the X and autosomes, 9 of 17 individuals have higher X than autosomal ancestry (*p* = 0.500, binomial test).

We find no statistical support for differences in X and autosomal ancestry, however, we cannot exclude low levels of sex-specific mating between early farmers and hunter-gatherers. Therefore, we evaluated the magnitude of differences in male and female contributions that would be consistent with observed X to autosomal ancestry ratios. We determined this range of sex bias values by simulating ancestry under a mechanistic admixture model including genetic drift and sampling at specified sample sizes (16,30,31) (Fig. 3A, Materials and Methods). Even for a small admixed population, the largest bias consistent with the observed X and autosomal ancestries is less than 1.2 males for every female, with a median over 1,000 simulations of 1.07.

**Fig. 3:**
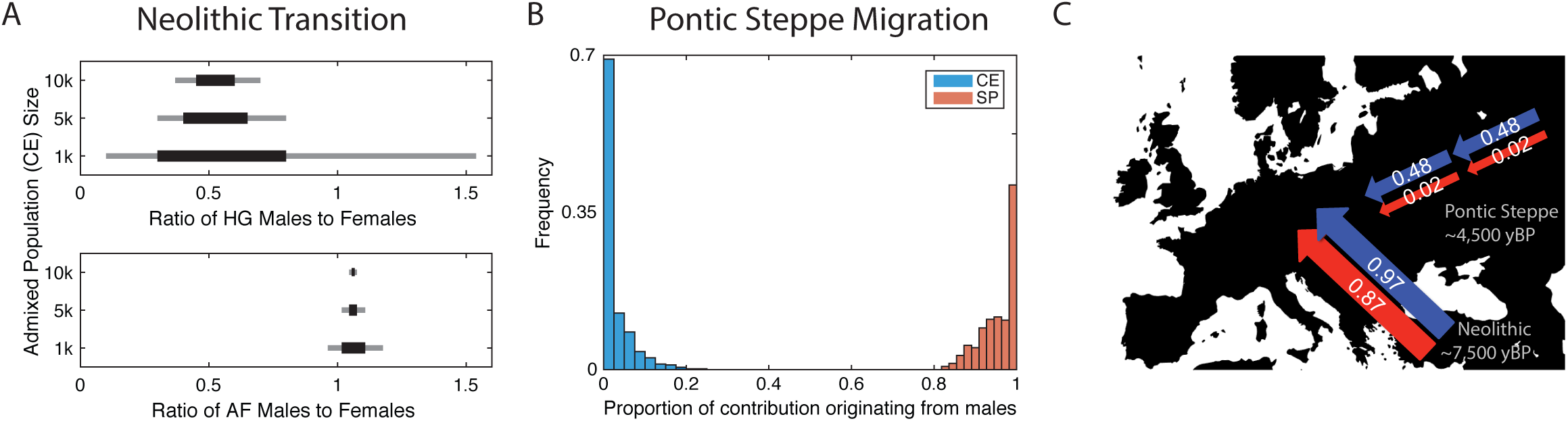
Estimated levels of sex bias during the Neolithic Transition and Pontic Steppe migration. (A) Neolithic Transition. The range of sex bias, measured as the ratio of males to females from a source population, that is consistent with the observed ratio of X and autosomal ancestries (Materials & Methods). Total contributions from the source population are specified based on autosomal ancestry as 0.913 from AF and 0.087 from HG. Lines indicate that the observed ratios of X to autosomal ancestry in our dataset were present in the middle 50% (black) or middle 80% (grey) of 1,000 simulated admixed populations for specified CE population sizes. (B) Pontic Steppe migration. Under a model of constant admixture over time, the fraction of the total contribution of genetic material originating from males for each source population; CE and SP. Contributions are estimated from the migration parameter sets that have the smallest 0.1% Euclidean distance between observed and model-calculated ancestries. (C) Schematic of sex-specific migrations during early and mid Holocene Europe. Female contributions in are in red, male contributions are in blue, estimated under a single pulse migration model from Anatolia, and under a constant migration model from the Pontic Steppe. The total contribution of each population is the average of female and male contributions from that source.

Consistent with the slightly larger X than autosomal ancestry observed for HG ancestry, under the simulation framework, we estimate a median of 1.91 females for every male from the HG population to early Neolithic Central Europeans. The signal of female bias in contributions from HG to CE might be caused by a male-biased inheritance structure in the new farming population. That is, it is possible that the migration from Anatolia involved substantial contributions from both men and women, but once in central Europe, a shift to patrilocality might have made absorption of local HG females easier than of HG males. However, the absolute difference between male and female contributions is small (~0.06). Correspondingly, differences in the numbers of female and male migrants would be small or are potentially a result of sampling.

Considering the these analyses together, we find no statistical support for a male-biased migration from Anatolia with a small range of possible sex bias values consistent with the data, and a potential signal of female-biased contributions from HG to CE.

We next considered female and male migration histories during the late Neolithic/Bronze Age migration from the Pontic-Caspian Steppe (Fig. 1). In contrast with the early Neolithic expansion from Anatolia, we find a strikingly lower distribution of SP ancestry on the X chromosome than the autosomes (Fig. 2, Fig. S1, in accordance with *F*_*ST*_ results), suggesting extreme male-biased migration from SP during the Late Neolithic/Bronze Age migration from the Pontic-Caspian Steppe. Using a similar approach as that employed for the early Neolithic migration event, the ratio of mean X-chromosomal SP ancestry to mean autosomal SP ancestry in late Neolithic and Bronze Age Europeans (BA) is 0.366/0.618 = 0.592. The ratio of mean X CE ancestry to mean autosomal CE ancestry in the BA population is 0.634/0.382 = 1.66. Of 16 admixed BA individuals, 12 have more SP ancestry on the autosomes than the X chromosome (binomial test, *p* = 0.038). Similarly, the distribution of p-values of the Wilcoxon sign-rank test comparing the estimated X-chromosomal ancestries to the autosomal ancestries in each of 100 resamples of autosomal SNPs is highly skewed toward zero, with a median of *p* = 0.02 (Materials & Methods, Fig. S2).

To interpret the values of sex-specific admixture that can produce the observed ratio of X-to-autosomal SP ancestry of about 0.6, we considered four models for the admixture process. The first is a single admixture event, in which an SP population quickly mixes with central European farmers, with no further migration from either population to the admixed BA population. Under this model, however, the level of sex bias is too high to have been produced by a single admixture event; no solution for the female and male migration rates exists within the possible admixture contribution range from 0 to 1 (Materials and Methods). In other words, in a pulse migration and admixture scenario in a single generation, even a male-only migration event is not extreme enough to generate the observed extreme X-to-autosome bias in the data. Ongoing male migration from the steppe over multiple generations is therefore required to explain observed patterns of X and autosomal ancestry.

We therefore considered a model of constant contributions over time from the SP population and early Neolithic farmers (CE). We follow the method of (16), comparing expected X and autosomal ancestry (16, eqs. 19,20; and 30, eq. 31) to observed ancestry in our data over a grid of possible parameter values. We present results from the 0.1% of parameter sets closest to observed data using a Euclidean distance between model-based and observed population mean ancestries on the X and autosomes (Materials and Methods). Other cutoffs (0.5, 1, 5%) showed similar trends.

Figure S3 plots the range of sex-specific contributions from the SP and CE populations that produce estimates close to those observed in the BA population. Males from the steppe and central European females show substantial ongoing migration, with continuing admixture rates of almost ½. That is, almost half of the male parents in each generation of BA individuals are new migrants from the SP population. Females from the steppe and early Neolithic European males, however, are estimated to have contributed negligibly to the BA population. Figure 3B plots the proportional contribution of males from each source population, with a median of about 94% of SP ancestry in the BA population coming from male SP migrants, and all local CE ancestry originating in CE females. This result corresponds to approximately 14 male migrants for every female migrant from the steppe contributing to the ancestry of the BA population. Considering the smallest 0.5%, 1% and 5% of Euclidean distances instead, this ratio is about 8.5, 7.5, and 5.1, respectively, males per female migrating from the steppe.

The signature of X-chromosomal to autosomal ancestry is driven by the last few generations of admixture. Testing other models of time-dependent admixture, with the contributions from one or both of the source populations increasing or decreasing over time, we find that the data fit model-based estimates approximately equally well when the admixture contributions at the last few generations are similar to those estimated from a constant admixture model (Materials and Methods).

The signal of male-biased contributions from SP and to BA over time is consistent with an admixture scenario in which a massive male-biased migration from the steppe initially looks to local European farmer females for wives, and with a paternal mode of inheritance, the BA population disproportionately absorbs females from local ‘unadmixed’ farmers. Admixture from the steppe population continues over time, though mainly men migrate, perhaps expanding using the male-dominated modes of horses and chariots (23,26).

Overall, the model-based ancestry results show remarkable similarity to our original comparisons of relative genetic drift on the X versus autosomes using a measure of genetic differentiation, *F*_*ST*_ (Table 1). Combining observations from both migrations, a picture of sex-specific migrations in central European prehistory emerges (Fig. 3C).

Owing to the large ancestry contribution and lack of sex-biased admixture, the massive cultural change that accompanied the shift to agriculture is consistent with a large-scale migration of an entire subset of a population, perhaps families, and a slower rate of spread. Minimal differences in sex-specific migration and the high overall AF ancestry in CE individuals support this scenario. This result suggests that the residence and descent rules were not determining factors in sex-specific migration, despite the probable patrilocality of the migrating AF population (9,22). The lack of sex bias is in fact consistent with previous indications of sex bias during the Neolithic based on mtDNA diversity. Earlier work focused on measures of diversity rather than ancestry, which will track the effective population size more than admixture. Therefore, earlier single-locus studies are likely seeing the signal of patrilocality rather than the migration process from Anatolia (19).

In contrast, our results, combined with the archeological evidence, suggest that the rapid migration from the Pontic Steppe was strongly male-biased, potentially via newly domesticated horses in multiple waves (23,24,26). Such differences in sex-specific migration patterns are suggestive of fundamentally different types of interactions between invading and local populations during the two migration events. Our results demonstrate the power for inferring important processes in human prehistory by analyzing the X chromosome and the autosomes jointly.

## Materials and Methods

### Genetic samples and populations

We analyzed published (6) ancient samples that have been genotyped for a set of 1,240,000 SNPs, including 49,711 on the X chromosome. Under notation from (6), for the early Neolithic migration from Anatolia, we considered individuals from the CEM population label for ‘selection label 2’; for the Late Neolithic/Bronze Age migrations from the Pontic Steppe, we considered individuals with ‘archeological culture’ label Central_LNBA. These subsets of the data geographically restrict analyses to Central Europeans, decreasing potential variation from spatial variation within Europe. Additionally, while the samples each span approximately one thousand years, the small correlations (< 0.1) between X or autosomal ancestry and calibrated dates are not statistically significant.

Additional genomic filtering and analyses were done in PLINK v1.90 (32). We removed the pseudoautosomal region of the X chromosome, and removed SNPs with pairwise correlation greater than 0.4 using the command ‘indep-pairwise 200 25 0.4’ following previous ancient DNA studies (4,6). We considered admixed individuals with at least 1,000 SNPs on the X chromosome. Tables S1 and S2 show the individuals used in analyses and their population classifications. More information on the samples is available in (6).

### Sex-biased genetic differentiation

As a first line of evidence for the sex-specific relationships between the two sets of migrating and admixed populations, AF-CE and SP-BA, we compared genetic differentiation on the X versus autosomes (Table 1). We followed the method of (27,28), computing the statistic *Q*, which measures relative genetic drift between the X and autosomes, 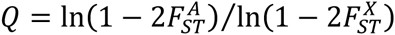. We calculated 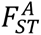 and 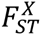 in Plink v1.9, using a ratio of averages approach to combine SNPs and (33, eq. 10)’s estimator.

Values of *Q* are suggestive, however, as deviations from ¾ can also be produced by population histories with population size or migration changes even in the absence of sex bias (34,35). Additionally, *Q* lacks a clear framework for quantitative interpretation. Therefore, we used a mechanistic admixture model comparing ancestry on the X chromosome and autosomes to infer sex-specific admixture rates and compare potential migration models.

### Estimating ancestry components

Evidence of admixture and migration events in the population history of Central Europeans, as well as current best proxy populations for their sources, has been extensively presented in other studies (1–6). Therefore, we assumed these migration events occurred, and used the best samples/populations currently available as representatives of relatives of the admixed populations. A schematic of the migration events is in Fig. 1, with estimated ancestry components in Fig. 2. Results by individual are presented in Tables S1, S3 and S4.

We estimated ancestry components of the two admixed populations, CE and BA. For the early Neolithic transition to agriculture, we assumed two ancestry components (*K*=2), with Anatolian Neolithic farmers (AF) and European hunter-gatherer (HG) source populations. For the later migration from the steppe, we assumed three ancestry components (*K*=3), with contributions from the Pontic Steppe (SP) represented by the Yamnaya Samara population, as well as contributions from AF and HG populations.

Multiple methods exist to infer individual ancestry proportions. We considered two of the most common clustering methods: *Admixture* (29) v1.3, a maximum likelihood method, and *Structure* (36) v2.3, a Bayesian algorithm. Both methods rely on a similar underlying model, with different estimation techniques. In each program, we tested supervised and unsupervised clustering and compared individual ancestry estimates and population-level summary statistic estimates between methods. Results are summarized in Tables S3 and S4. We did not use f-statistic based ancestry estimates for X-chromosomal ancestry as they depend on effective population size, and therefore cannot be compared between the X chromosome and autosomes, which have different effective sizes (37).

Estimated individual ancestry from supervised and unsupervised clustering in *Admixture* are highly concordant for the autosomes for both migration events, and for the X chromosome for the Neolithic migration scenario. For X-chromosomal ancestry estimated for the steppe migration, however, reference individuals do not emerge as clusters in unsupervised *Admixture*, therefore results cannot be used in this framework. For both the X chromosome and autosomes, unsupervised *Structure* estimates are similar to those from *Admixture*. Perhaps surprisingly, supervised *Structure* produces different estimates for both individual and population-level ancestry estimates, but as we show below, this is likely an effect of a small sample size, where increasing the sample size for supervised *Structure* leads to results similar to *Admixture* and unsupervised *Structure* for an example case with high coverage data. In all *Structure* analyses, we use 10,000 burn-in iterations, followed by 50,000 iterations.

As our inference methods rely on the mean ancestry in the population (and are supported by the variance), we focus on comparisons of these summary statistics. The population mean and variance of ancestry for supervised *Structure,* supervised and unsupervised *Admixture* are highly concordant (no data for BA X chromosome in unsupervised *Admixture*). However, the mean ancestry estimated in supervised *Structure* is qualitatively different, and the variance is substantially lower. That is, downstream analyses using the first three methods would produce similar estimates of the levels of sex bias during the two migrations, while inference from supervised *Structure* would be qualitatively different, though notably, still in the same direction we observe for both migration events.

*Comparing ancestry estimation methods using HapMap data:* As three methods produce concordant results, with only supervised *Structure* differing, we investigated various factors, particularly sample size, that can cause the deviation in supervised *Structure* results. We investigated the effect of sample size by reproducing the analysis on a larger dataset from the HapMap Phase 3 Project (38). We considered the recent admixture between Africans and Europeans, using YRI and CEU as reference populations and estimating ancestry in ASW individuals (Figure S4). To examine the impact of sample size on accuracy of ancestry inference, we down sampled the reference populations considering sample sizes ranging from 4 diploid individuals in each reference population to 112 (the maximum number of CEU individuals). The set of individuals in smaller reference panels are subsets of the larger panels.

For each sample size of reference populations, we estimated ancestry in 16 ASW individuals using each of the four methods (supervised and unsupervised settings in each *Admixture* and *Structure*), based on the sample size of BA. Figure S4 plots the mean and variance of ancestry in the ASW population for each sample size and method. Each point is based on 10 replicates, first averaging ancestry by individual. For computational speed, we estimated ancestry from 15,000 randomly drawn autosomal SNPs, after LD pruning using the same method as for the ancient DNA (see Genetic samples and populations). For each replicate, we resampled 15,000 SNPs and use a new seed. As in our data, supervised *Structure* is an outlier in its behavior. Indeed, as the ancient samples are largely haploid, the corresponding sample sizes most representative are in the range of four to ten diploid CEU/YRI individuals. Particularly in this range of the plot, the mean and variance of ancestry estimated using supervised *Structure* differ greatly from those using all other methods, and are further from corresponding estimates at large sample sizes under all methods (Figure S4). As the sample sizes increase for the reference populations, the results from supervised *Structure* approach the results of the three other methods (Figure S4). We conclude that the differing finding using supervised *Structure* for the ancient individuals is due to sensitivity to low sample sizes for the supervised *Structure* algorithm. Therefore, we use estimates from supervised *Admixture* for inference in the main text, and reiterate that inference using unsupervised *Structure* would be highly similar because of the similar mean and variance of ancestry estimated. Ancestry for each individual is presented as the average estimated individual ancestry using ten independent seeds, considering the X chromosome and autosomes separately.

To compare the X chromosome to the autosomes, we estimated autosomal ancestry on 100 sets of SNPs resampled from the autosomes. For each of the two migration events, we resampled autosomal ancestry to match the number of SNPs used in X-chromosomal analyses. For the Neolithic transition, the number of SNPs was 3,763. For the steppe migration, the number of SNPs was 4,605. We also down-sampled X-chromosomal SNPs for BA individuals to 3,763 ten times to compare estimates of ancestry between the two migration events. Ancestry estimates based on the down-sampled data were within 5% of original full data.

To test if the ancestry estimates are stable over the choice of individuals in the source populations, we tested multiple subsets of source population individuals (Table S2): 1) all individuals from (6) for the respective categories: using original population descriptions, Anatolians, Western + Scandinavian HGs, and Yamnaya Samara, 2) the subset of individuals whose genetic population assignment matches their known cultural association in an average of 10 independent unsupervised admixture runs for both the X and the autosomes.

For the first event, the Neolithic transition, the estimated ancestry components are roughly constant with varying choice of individuals. For the migration from the steppe, however, we see a range of values for estimated ancestries over different seeds, suggesting variation in the likelihood surface. The qualitative results are consistent through all analyses. The population means for the X chromosome and autosomes range from 0.27 to 0.44, and 0.54 to 0.73, respectively. The ratio of X to autosomal ancestry for a given seed varies between 0.38 and 0.61. While the magnitude of ancestry estimates varies, the signal of substantial sex bias based on the ratio of X to autosomal ancestry is seen for all scenarios.

For all analyses, we used ancestry estimated from the mean per individual of X-chromosomal estimates over the ten seeds. Autosomal ancestry is estimated as the median of 100 estimates from resampled SNP sets, which is in the lower range of autosomal estimates. For the steppe migration, this leads to a ratio of mean X-chromosomal to mean autosomal ancestry of 0.59, which is on the conservative (closer to 1) end of the range of estimates.

### Statistical significance of X and autosomal differences

We tested for statistical significance of the difference between the population means of X and autosomal ancestry within the admixed Neolithic population using the Wilcoxon sign-rank test. We did 100 comparisons of the distribution of ancestry on the X chromosome within the population to the distribution of autosomal ancestry estimated using each resample of *M* SNPs, where *M* is the number of X-chromosomal SNPs for the associated population (see Estimating Ancestry Components).

The Wilcoxon sign-rank test is a non-parametric paired difference test. For a statistically significant difference in the within population distribution of X and autosomal ancestry, one would expect an excess of small p-values. Rather, for the Neolithic transition, comparing X and autosomal AF-related ancestry, the p-value distribution over the 100 calculations is approximately uniform (Fig. S2). Similarly, comparing X and autosomal ancestry when autosomal ancestry estimated from all SNPs together (*M*=331,515) the Wilcoxon sign-rank test is not significant (*p* = 0.493). In contrast, for the later migration from the Pontic Steppe, comparing the distribution of ancestry on the X chromosome to that estimated for the autosomes with all SNPs together (*M*=375,243), *p* = 0.002, and we see an excess of small p values for the comparisons to 100 resampled autosomal estimates (Fig. S2).

### Simulations to estimate range of sex bias during Neolithic Transition

For a constant admixed population of size *N*, with *N* ϵ {1,000; 5,000; 10,000}, we simulated the ancestry proportion of individuals in the admixed population recursively for 40 generations, or approximately 1,000 years, assuming a single admixture event followed by no further migration (Fig. 1A). For a generation time of ~25 years, this number of generations approximately corresponds to the difference in time between the onset of migration and the radiocarbon ages of sampled admixed individuals for each migration (6).

We set the total contributions from each population based on their autosomal ancestry levels (16,32,33), with HG as 0.087, and AF to be 0.913. Given this constant level of contributions from the two source populations, we then did 1,000 replicate simulations for different levels of specified sex bias. We define the level of sex bias as the ratio of male to female contributions from a given source population, *B*, considering 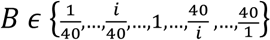.

Given the overall contribution from each source population, as well as a specified value of male to female contributions, *B,* the female and male contribution parameters can be exactly solved. That is, adapting eq. 1 from (33), for male contribution from population *α* given by *m*_*α*_, the probability of a randomly chosen individual in the first generation of the admixed population having a male parent from each source population is

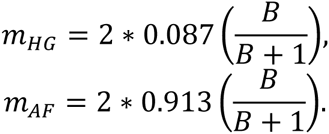

Similarly, the female contributions (*f*_*α*_) can be written as

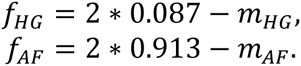

*Simulating autosomal ancestry:*for the first generation, *g* = 1, we randomly chose and matched *2N* parents, making *N* parental pairs. Each parent is drawn with probability given by their sex-specific contribution level. An individual’s autosomal ancestry is calculated as the average of its parent’s autosomal ancestries. Then, for *g* ≥ *2*, we calculated ancestry in *N* individuals at generation *g* by randomly choosing and pairing *2N* parents from the population in the previous generation, *g* − 1.

*Simulating X-chromosomal ancestry*: we followed the same procedure as for the autosomes, but instead considered separate populations of males and females, each with *N*/*2* individuals. For the female population, we generated *N* parental pairs by drawing an individual from each the male and female population from the previous generation. For the male population, we drew *N*/*2* mothers from the female population in the previous generation. Ancestry of females was calculated as the average of the parental ancestries, while ancestry for males is equal to the ancestry of the mother.

At *g* = 40, we randomly sampled 20 individuals, and calculated the mean autosomal ancestry and mean X-chromosomal ancestry in the sample. The mean X-chromosomal ancestry is calculated as a weighted mean of the female and male X-chromosomal ancestries, based on the proportion of females in the data set (75%). Figure 3A shows the values of sex bias, *B*, for which the observed X to autosomal ancestry ratio is within the middle 50% and 80% of ratios calculated from the 1,000 simulated populations with that level of specified sex bias.

The effect of drift on admixture fractions is larger in smaller populations (39); we therefore expect a larger possible range of sex bias values to produce values of X-to-autosomal ancestry ratios similar to those estimates from the data for smaller population sizes. Yet, even simulations with an admixed population size of 1,000 suggest less than 1.2 males migrating for every female from AF to CE.

### Admixture models for migration from the Steppe

We used recursive expressions for X and autosomal ancestry as a function of sex-specific admixture rates to interpret observed ancestry (16,31). We considered four general models of admixture over time: 1) single admixture event, with no further migration, 2) constant migration over time, 3) increasing migration from SP over time, 4) decreasing migration from SP over time.

First, we considered a single pulse admixture event, analogous to that used for the early Neolithic Migration from Anatolia. For mean autosomal ancestry within the admixed population of 0.618 and mean X-chromosomal ancestry of 0.366, under the model of a single admixture event with no further migration, we used eqs. 22-23 from (16) to write X and autosomal ancestries as a function of sex-specific contribution parameters. We have

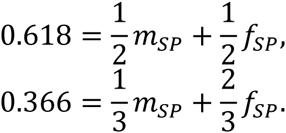

However, no solution exists within the bounds on migration contributions of *m*_*SP*_, *f*_*SP*_ ϵ [0,1].

We next considered constant admixture over time. Assuming *g* = 40, we computed the mean female and male X-chromosomal and autosomal admixture components (16, eqs. 5, 17,18) on a grid of possible sex-specific contribution parameter values *m*_*SP*_, *f*_*SP*_, *m*_*CE*_, *f*_*CE*_, ϵ [0,1], in 0.02 increments. We fixed initial values to be equal and without sex bias. Mean ancestry levels approach a limit around 15 generations, therefore initial conditions do not significantly impact final ancestries (16,30,31).

As the number of males in both admixed populations is small, mean sample ancestry estimates may not be representative of the population mean. Therefore, we followed eq. 25 from (16), calculating a pooled female and male Euclidean distance between model-based ancestry calculations and observed ancestry estimates. Figure 2 present results based on the smallest 0.1% of Euclidean distances on the grid, with estimated sex bias values for other cutoff in the text.

For time-dependent admixture rates, admixture per generation is calculated as a linear function of the number of generations spanning 0 to the contribution specified by that point on the grid corresponding to the constant admixture scenario. We used the recursive expressions from (16, eqs. 5,17,18) to calculate mean X and autosomal ancestry for each point on the grid.

These scenarios however are one of many that are possible, and further work is needed to describe the spatiotemporal variation in admixture during both the Neolithic migration and the later steppe migration. While spatially and temporal resolution will refine admixture models, the signals of sex-specific admixture during the prehistory of Central Europe will persist. Similarly, other processes may also differentially affect the X chromosome and autosomes, including recombination, mutation and selection, but these forces are unlikely to have a large impact on the chromosome-wide ancestry-based summary statistics we base analyses on over the short time scales considered.

### Variance in ancestry

Our analyses focus on comparisons of mean X-chromosomal and autosomal ancestry. However the variance can also be informative about the admixture history (30,31). The variance in ancestry with the admixed Neolithic individuals is quite low (0.013 for the X chromosome and 0.005 for the autosomes), with a higher variance in the admixed BA population (0.102 for the X chromosome and 0.039 for the autosomes). Larger X than autosomal variance is expected owing to the difference in the number of chromosomes inherited per generation. The higher variance in ancestry across individuals associated with the Pontic Steppe migration is consistent with recent or ongoing migration within the past few generations, particularly as sex bias would decrease the variance (31). Additionally, with recent or ongoing male-biased migration, one would expect lower Steppe ancestry on X chromosomes in admixed males than in admixed females, as females receive an X chromosome from their fathers. The mean X-chromosomal ancestry in BA males is roughly half that of BA females, though the difference is not statistically significant with only four individuals. While consistent with inferences from mean ancestry components, strong conclusions cannot be drawn from the variance or differences in male and female ancestry given the current sample sizes.

## Acknowledgements

We thank Jonathan Kang for bioinformatics support with the HapMap data. We acknowledge support from NSF Graduate Research and ARCS fellowships to AG, Wenner-Gren Foundation fellowship to TG, NSF grant BCS 1515127 to NAR, ERC grant #311413 to MJ, as well as an NSF-Swedish Research Council GROW award for AG to visit Uppsala University. AG and MJ conceived the project; MJ headed the project; AG, TG, NAR, and MJ designed the project; AG analyzed the data and performed the simulations with support from TG; and AG, TG, NAR, and MJ wrote the paper.

---

**Figure S1:**
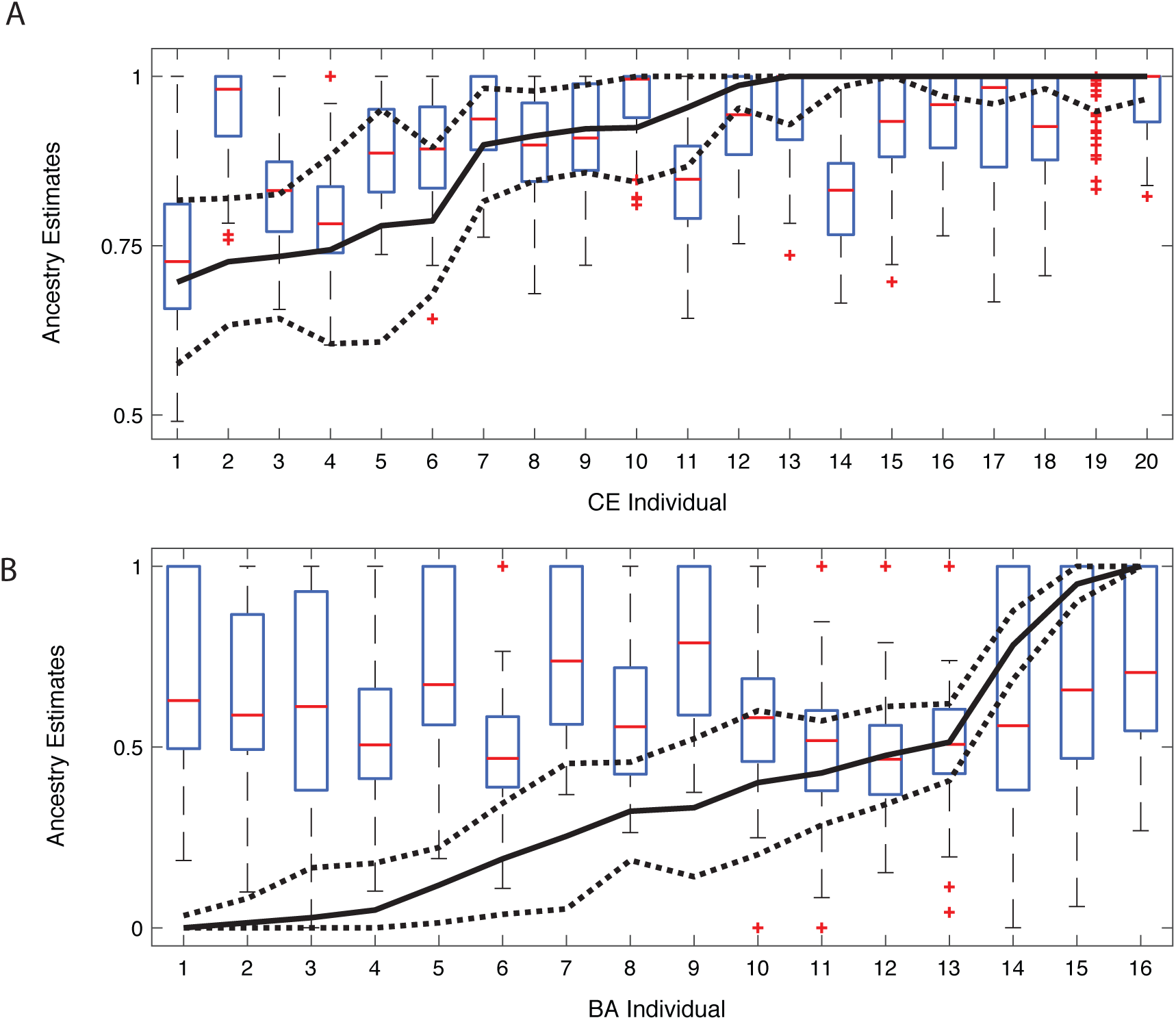
Resampling SNPs to estimate autosomal ancestry. Each boxplot represents the range of estimated autosomal ancestries over 100 resampled sets of randomly drawn autsomal SNPs to match the number of X-chromosomal SNPs. The lines correspond to estimated X-chromosomal ancestry by individual, with dotted lines marking within one standard error. (A) The Neolithic Transition. The number of SNPs is 3,763. The distribution of X and autosomal ancestry largely overlap in most individuals. (B) The Pontic Steppe migration. The number of SNPs is 4,605. Ancestry on the X is either lower or similar for all individuals. For both panels, individuals are presented in the same order as Fig. 2.

**Figure S2:**
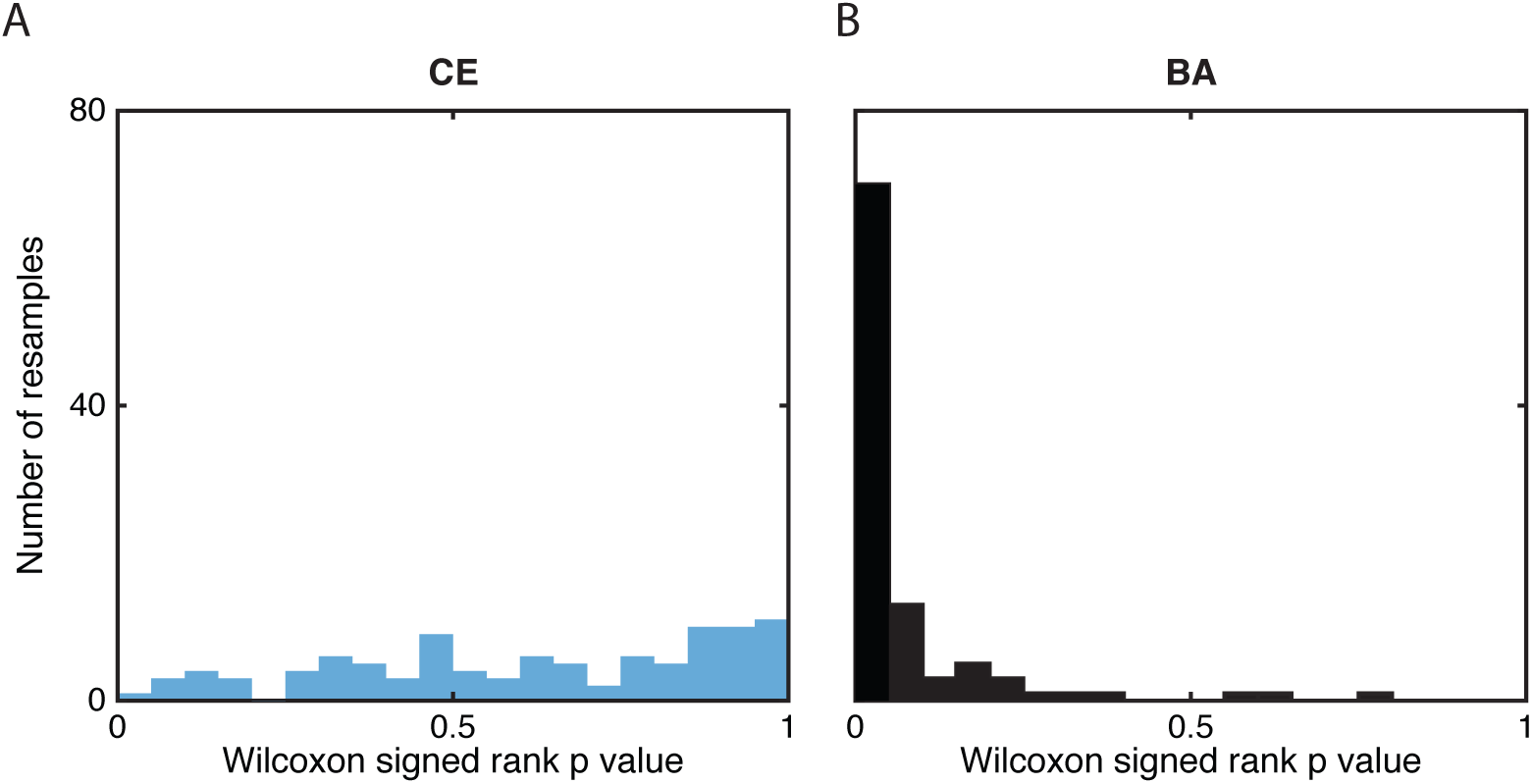
Histogram of p values for the Wilcoxon sign-rank test. In each admixed population, the comparison of the distribution of ancestry on the X chromsome to the 100 autosomal ancestry distributions for (A) the CE population, and (B) the BA population.

**Figure S3:**
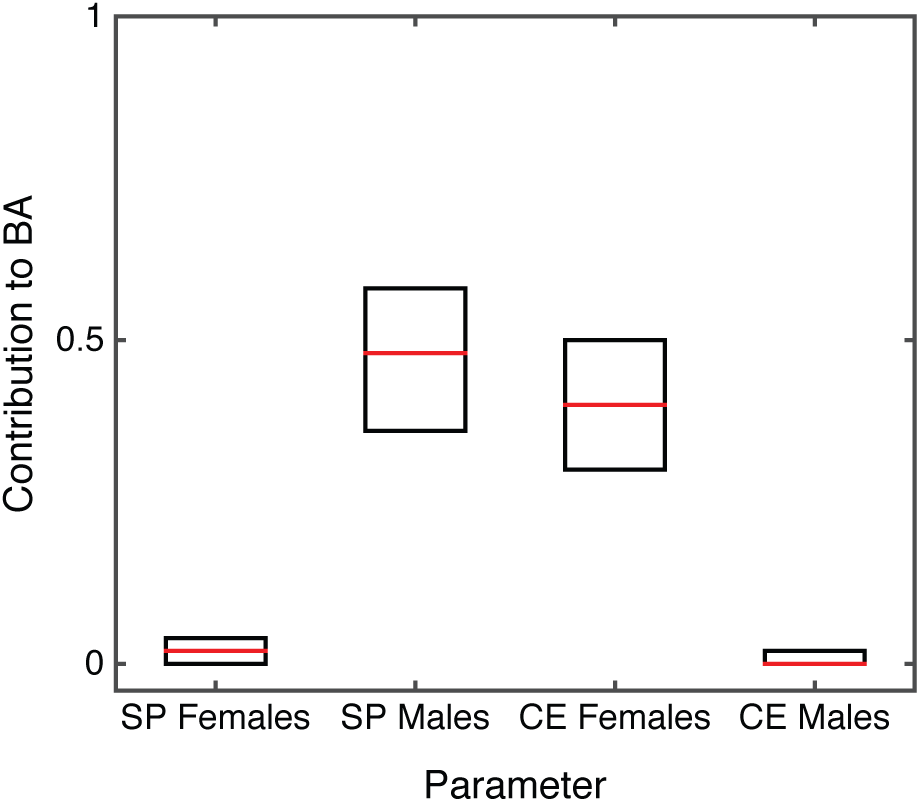
Estimated sex-specific contributions from Pontic Steppe (SP) and early Central Europeans (CE) to LNBA Europeans (BA). Under a model of constant contributions over time, each box represents the middle fifty percent of parameter sets for the smallest 0.1% of Euclidean distances between the model-predicted and observed X and autosomal ancestry from a grid of possible parameter values. Red line is the median of plotted values.

**Figure S4:**
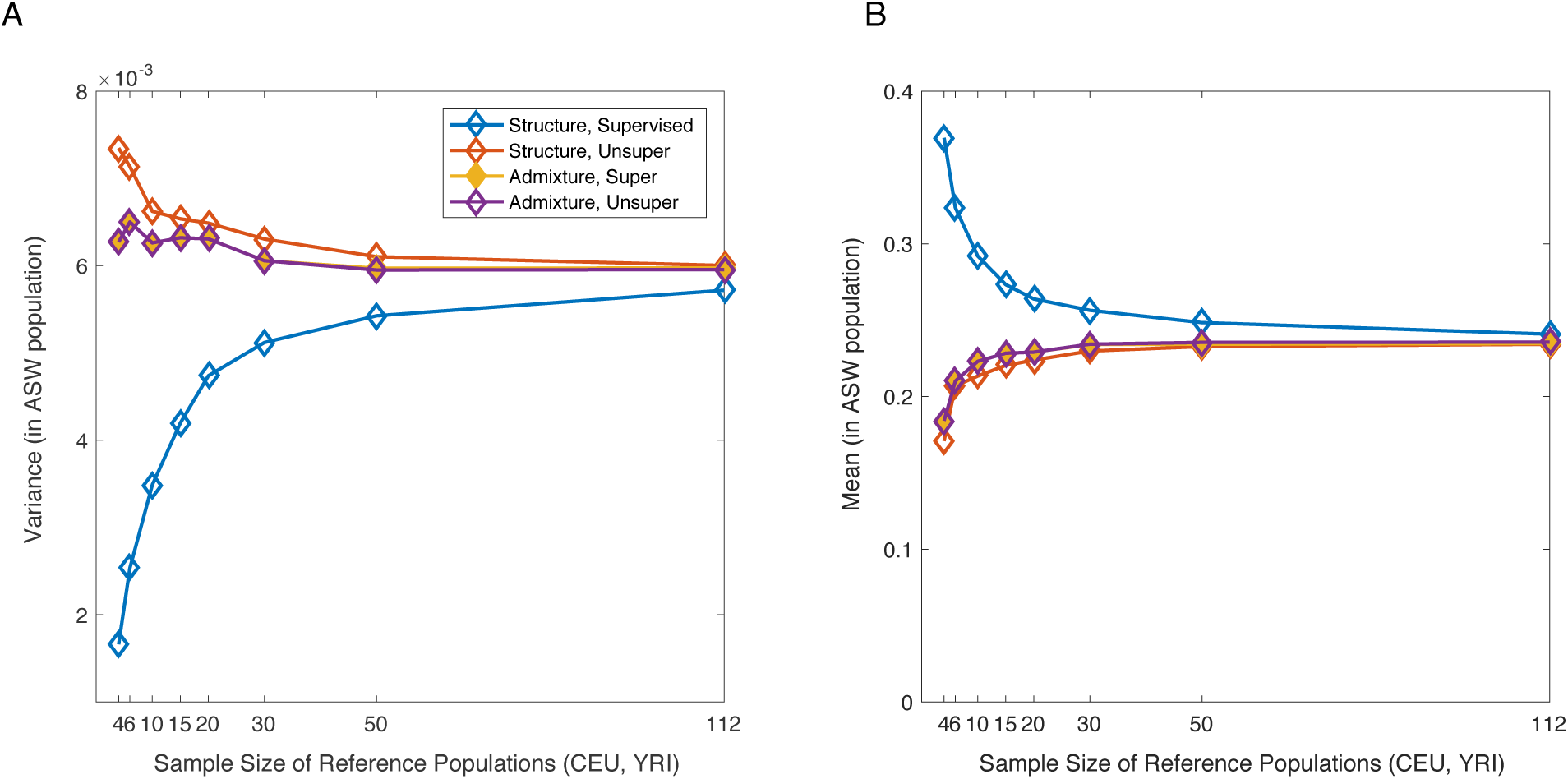
Comparing ancestry estimation methods using 1000 Genomes data. The (A) variance and (B) mean of ancestry in the ASW (African American) population by sample size for four clustering methods. Each point is based on 6 replicates, first averaging ancestry by individual. For each replicate, we resample 15,000 SNPs and use a new seed. Supervised clustering in *Structure* produces qualitatively different results than the other methods for small sample sizes.

**Table S1:** Individual ancestry in admixed populations. Ancestry is estimated in ADMIXTURE v1.3 using supervised clustering, results presented are the component clustering with the migrating population. For the Neolithic Transition, that is AF ancestry, for Pontic Steppe migration, that is SP ancestry. NT— Neolithic Transition, PSM—Pontic Steppe Migration.

| Migration Event | Individual | # SNPs (X) | # SNPs (Autosomes) | Ancestry, X chromosome | Ancestry, Autosomes (All SNPs) | Ancestry, Autosomes (Median of 100 sets) |
| --- | --- | --- | --- | --- | --- | --- |
| NT | I0172 | 3,348 | 296,827 | 0.74 | 0.80 | 0.78 |
| NT | I0560 | 1,293 | 125,647 | 0.70 | 0.74 | 0.73 |
| NT | I1497 | 2,945 | 282,608 | 1.00 | 0.85 | 0.83 |
| NT | I1495 | 2,595 | 282,090 | 0.78 | 0.89 | 0.89 |
| NT | I1496 | 2,504 | 277,979 | 1.00 | 1.00 | 1.00 |
| NT | I1498 | 2,944 | 281,643 | 0.79 | 0.89 | 0.89 |
| NT | I1499 | 2,979 | 282,181 | 0.95 | 0.86 | 0.85 |
| NT | I1500 | 2,540 | 282,155 | 0.90 | 0.95 | 0.94 |
| NT | I1505 | 2,835 | 271,675 | 0.92 | 0.91 | 0.91 |
| NT | I1506 | 2,553 | 250,572 | 0.91 | 0.91 | 0.90 |
| NT | I1508 | 2,118 | 202,813 | 1.00 | 1.00 | 1.00 |
| NT | Iceman | 3,563 | 329,264 | 0.73 | 0.82 | 0.83 |
| NT | I0022 | 1,226 | 127,899 | 1.00 | 1.00 | 0.98 |
| NT | I0025 | 2,838 | 276,166 | 1.00 | 0.94 | 0.93 |
| NT | I0026 | 3,002 | 287,954 | 0.92 | 0.99 | 1.00 |
| NT | I0046 | 3,008 | 281,010 | 0.99 | 0.98 | 0.94 |
| NT | I0054 | 3,269 | 305,318 | 1.00 | 0.95 | 0.96 |
| NT | I0100 | 3,159 | 309,476 | 0.73 | 0.96 | 0.98 |
| NT | I0659 | 1,186 | 181,302 | 1.00 | 0.99 | 1.00 |
| NT | I1550 | 1,827 | 178,442 | 1.00 | 0.96 | 0.93 |
| **Mean Ancestry:** |  |  |  | **0.903** | **0.919** | **0.913** |
| PSM | I0047 | 3,377 | 295,581 | 0.51 | 0.55 | 0.511 |
| PSM | I0049 | 2,314 | 190,443 | 0.33 | 1.00 | 1.000 |
| PSM | I0059 | 3,124 | 273,321 | 0.43 | 0.54 | 0.519 |
| PSM | I0099 | 3,081 | 339,040 | 0.19 | 0.54 | 0.479 |
| PSM | I0103 | 4,273 | 358,176 | 1.00 | 1.00 | 0.732 |
| PSM | I0104 | 3,223 | 336,630 | 0.25 | 1.00 | 0.789 |
| PSM | I0115 | 1,513 | 130,840 | 0.03 | 0.66 | 0.599 |
| PSM | I0116 | 1,353 | 229,560 | 0.40 | 0.63 | 0.589 |
| PSM | I0117 | 3,520 | 307,312 | 0.05 | 0.56 | 0.516 |
| PSM | I0118 | 4,205 | 364,382 | 0.48 | 0.54 | 0.493 |
| PSM | I0164 | 3,278 | 301,603 | 0.32 | 0.62 | 0.561 |
| PSM | I0803 | 1,241 | 84,086 | 0.78 | 0.68 | 0.573 |
| PSM | I1532 | 1,248 | 171,544 | 0.00 | 1.00 | 0.625 |
| PSM | RISE00 | 2,538 | 221,453 | 0.95 | 0.68 | 0.642 |
| PSM | RISE150 | 2,383 | 248,413 | 0.12 | 1.00 | 0.674 |
| PSM | RISE577 | 2,857 | 254,460 | 0.01 | 0.61 | 0.582 |
| **Mean Ancestry:** |  |  |  | **0.366** | **0.726** | **0.618** |

**Table S2:** Source population individuals. Ancestry is estimated in *Admixture* using supervised clustering, results presented are the component clustering with the migrating population. Individuals in italicized rows were not included in main text analyses (Materials & Methods). HG—Hunter gatherer, AF—Anatolian Farmer, SP—Pontic-Caspian Steppe.

| Population | Individual | # SNPs<br>(X) | # SNPs<br>(Autosomes) |
| --- | --- | --- | --- |
| HG | I0011 | 2,597 | 238,300 |
| HG | I0012 | 2,248 | 288,233 |
| HG | I0013 | 294 | 114,115 |
| HG | I0014 | 2,847 | 277,370 |
| HG | I0015 | 1,394 | 224,569 |
| HG | I0017 | 2,826 | 326,861 |
| HG | I0585 | 3,315 | 331,453 |
| HG | I1507 | 2,881 | 304,651 |
| HG | Loschbour | 4,483 | 370,431 |
| AF | I0707 | 3,928 | 347,728 |
| AF | I0708 | 3,120 | 339,272 |
| AF | I0709 | 3,196 | 344,000 |
| *AF* | *I0723* | *1,069* | *158,218* |
| *AF* | *I0724* | *101* | *20,982* |
| *AF* | *I0726* | *910* | *84,012* |
| *AF* | *I0727* | *95* | *16,374* |
| AF | I0736 | 3,090 | 293,649 |
| AF | I0744 | 2,665 | 316,049 |
| AF | I0745 | 3,290 | 346,243 |
| AF | I0746 | 3,310 | 347,428 |
| AF | I1096 | 2,658 | 292,505 |
| AF | I1097 | 2,532 | 289,766 |
| AF | I1098 | 3,401 | 302,251 |
| AF | I1099 | 1,579 | 215,557 |
| AF | I1100 | 1,338 | 121,755 |
| AF | I1101 | 2,176 | 259,865 |
| AF | I1102 | 1,087 | 168,149 |
| AF | I1103 | 1,900 | 239,704 |
| AF | I1579 | 3,525 | 306,814 |
| AF | I1580 | 3,853 | 332,503 |
| AF | I1581 | 3,578 | 308,641 |
| AF | I1583 | 3,665 | 349,463 |
| AF | I1585 | 3,621 | 309,674 |
| SP | I0231 | 3,695 | 361,283 |
| SP | I0357 | 2,298 | 189,734 |
| SP | I0370 | 2,047 | 254,721 |
| SP | I0429 | 1,600 | 224,264 |
| SP | I0438 | 1,353 | 206,462 |
| *SP* | *I0439* | *600* | *96,300* |
| SP | I0441 | 328 | 37,237 |
| SP | I0443 | 3,344 | 347,381 |
| SP | I0444 | 1,345 | 197,552 |

**Table S3:** Replication of individual autosomal ancestry in admixed populations. Ancestry is estimated in *Admixture* using unsupervised clustering, and in *Structure* using both supervised and unsupervised clustering. Results presented are the component clustering with the migrating population. For the Neolithic Transition, that is AF ancestry, for Pontic Steppe migration, that is SP ancestry. Results are the mean by individual of ten independent runs. We use ten different subsamples of 25,000 SNPs because of *Structure* run times. NT—Neolithic Transition, PSM—Pontic Steppe Migration.

| Migration Event | Individual | Ancestry, Admixture Unsupervised | Ancestry, Structure Unsupervised | Ancestry, Structure Supervised |
| --- | --- | --- | --- | --- |
| NT | I0172 | 0.83 | 0.83 | 0.75 |
| NT | I0560 | 0.77 | 0.86 | 0.76 |
| NT | I1497 | 0.86 | 0.89 | 0.78 |
| NT | I1495 | 0.90 | 0.86 | 0.77 |
| NT | I1496 | 1.00 | 1.00 | 0.83 |
| NT | I1498 | 0.90 | 0.96 | 0.79 |
| NT | I1499 | 0.87 | 0.86 | 0.78 |
| NT | I1500 | 0.95 | 1.00 | 0.83 |
| NT | I1505 | 0.92 | 0.93 | 0.79 |
| NT | I1506 | 0.92 | 0.91 | 0.79 |
| NT | I1508 | 1.00 | 1.00 | 0.85 |
| NT | Iceman | 0.85 | 0.85 | 0.76 |
| NT | I0022 | 1.00 | 1.00 | 0.84 |
| NT | I0025 | 0.95 | 0.99 | 0.80 |
| NT | I0026 | 0.99 | 1.00 | 0.82 |
| NT | I0046 | 0.98 | 0.98 | 0.80 |
| NT | I0054 | 0.95 | 0.95 | 0.79 |
| NT | I0100 | 0.97 | 1.00 | 0.82 |
| NT | I0659 | 0.99 | 1.00 | 0.82 |
| NT | I1550 | 0.96 | 0.96 | 0.80 |
| **Mean Ancestry:** |  | **0.928** | **0.941** | **0.798** |
| PSM | I0047 | 0.59 | 0.65 | 0.24 |
| PSM | I0049 | 1.00 | 0.99 | 0.26 |
| PSM | I0059 | 0.59 | 0.58 | 0.23 |
| PSM | I0099 | 0.59 | 0.67 | 0.23 |
| PSM | I0103 | 1.00 | 0.92 | 0.27 |
| PSM | I0104 | 1.00 | 0.79 | 0.28 |
| PSM | I0115 | 0.70 | 0.69 | 0.26 |
| PSM | I0116 | 0.68 | 0.77 | 0.22 |
| PSM | I0117 | 0.61 | 0.63 | 0.22 |
| PSM | I0118 | 0.58 | 0.68 | 0.22 |
| PSM | I0164 | 0.67 | 0.71 | 0.25 |
| PSM | I0803 | 0.75 | 0.69 | 0.26 |
| PSM | I1532 | 1.00 | 0.81 | 0.24 |
| PSM | RISE00 | 0.75 | 0.93 | 0.24 |
| PSM | RISE150 | 1.00 | 0.83 | 0.25 |
| PSM | RISE577 | 0.67 | 0.80 | 0.25 |
| **Mean Ancestry:** |  | **0.761** | **0.759** | **0.244** |

**Table S4:** Replication of individual X-chromosomal ancestry in admixed populations. Ancestry is estimated in *Admixture* using unsupervised clustering, and in *Structure* using both supervised and unsupervised clustering. Results presented are the component clustering with the migrating population. For the Neolithic Transition, that is AF ancestry, for Pontic Steppe migration, that is SP ancestry. Results are the mean by individual of ten independent runs. NT—Neolithic Transition, PSM—Pontic Steppe Migration.

| Migration Event | Individual | Ancestry, Admixture Unsupervised | Ancestry, Structure Unsupervised | Ancestry, Structure Supervised |
| --- | --- | --- | --- | --- |
| NT | I0172 | 0.72 | 0.86 | 0.72 |
| NT | I0560 | 0.72 | 0.85 | 0.71 |
| NT | I1497 | 1.00 | 1.00 | 0.80 |
| NT | I1495 | 0.73 | 0.93 | 0.74 |
| NT | I1496 | 1.00 | 1.00 | 0.85 |
| NT | I1498 | 0.78 | 0.83 | 0.72 |
| NT | I1499 | 0.91 | 0.98 | 0.80 |
| NT | I1500 | 0.90 | 0.94 | 0.74 |
| NT | I1505 | 0.90 | 0.94 | 0.76 |
| NT | I1506 | 0.89 | 0.97 | 0.78 |
| NT | I1508 | 1.00 | 1.00 | 0.85 |
| NT | Iceman | 0.73 | 0.77 | 0.70 |
| NT | I0022 | 1.00 | 1.00 | 0.86 |
| NT | I0025 | 0.99 | 1.00 | 0.78 |
| NT | I0026 | 0.91 | 0.97 | 0.75 |
| NT | I0046 | 0.98 | 1.00 | 0.77 |
| NT | I0054 | 1.00 | 1.00 | 0.79 |
| NT | I0100 | 0.72 | 0.73 | 0.70 |
| NT | I0659 | 1.00 | 1.00 | 0.81 |
| NT | I1550 | 1.00 | 1.00 | 0.80 |
| **Mean Ancestry:** |  | **0.894** | **0.937** | **0.772** |
| PSM | I0047 | - | 0.46 | 0.21 |
| PSM | I0049 | - | 0.46 | 0.20 |
| PSM | I0059 | - | 0.52 | 0.23 |
| PSM | I0099 | - | 0.18 | 0.16 |
| PSM | I0103 | - | 0.91 | 0.26 |
| PSM | I0104 | - | 0.28 | 0.18 |
| PSM | I0115 | - | 0.04 | 0.15 |
| PSM | I0116 | - | 0.63 | 0.20 |
| PSM | I0117 | - | 0.15 | 0.15 |
| PSM | I0118 | - | 0.53 | 0.21 |
| PSM | I0164 | - | 0.35 | 0.19 |
| PSM | I0803 | - | 0.66 | 0.22 |
| PSM | I1532 | - | 0.00 | 0.12 |
| PSM | RISE00 | - | 0.72 | 0.20 |
| PSM | RISE150 | - | 0.15 | 0.19 |
| PSM | RISE577 | - | 0.01 | 0.16 |
| **Mean Ancestry:** |  | **NA** | **0.378** | **0.189** |

